# A proteogenomic atlas of the human neural retina

**DOI:** 10.1101/2024.05.22.595273

**Authors:** Tabea V. Riepe, Merel Stemerdink, Renee Salz, Alfredo Dueñas Rey, Suzanne E. de Bruijn, Erica Boonen, Tomasz Z Tomkiewicz, Michael Kwint, Jolein Gloerich, Hans J.C.T. Wessels, Elfride De Baere, Filip van Nieuwerburgh, Sarah De Keulenaer, Barbara Ferrari, Stefano Ferrari, Frauke Coppieters, Frans P.M. Cremers, Erwin van Wyk, Susanne Roosing, Erik de Vrieze, Peter A.C. ‘t Hoen

## Abstract

The human neural retina is a complex tissue with abundant alternative splicing and more than 10% of genetic variants linked to inherited retinal diseases (IRDs) alter splicing. Traditional short-read RNA-sequencing methods have been used for understanding retina-specific splicing but have limitations in detailing transcript isoforms. To address this, we generated a proteogenomic atlas that combines PacBio long-read RNA-sequencing data with mass spectrometry and whole genome sequencing data of three healthy human neural retina samples. We identified nearly 60,000 transcript isoforms, of which approximately one-third are novel. Additionally, ten novel peptides confirmed novel transcript isoforms. For instance, we identified a novel *IMPDH1* isoform with a novel combination of known exons that is supported by peptide evidence. Our research underscores the potential of in-depth tissue-specific transcriptomic analysis to enhance our grasp of tissue-specific alternative splicing. The data underlying the proteogenomic atlas are available via EGA with identifier EGAD50000000101, via ProteomeXchange with identifier PXD045187, and accessible through the UCSC genome browser.

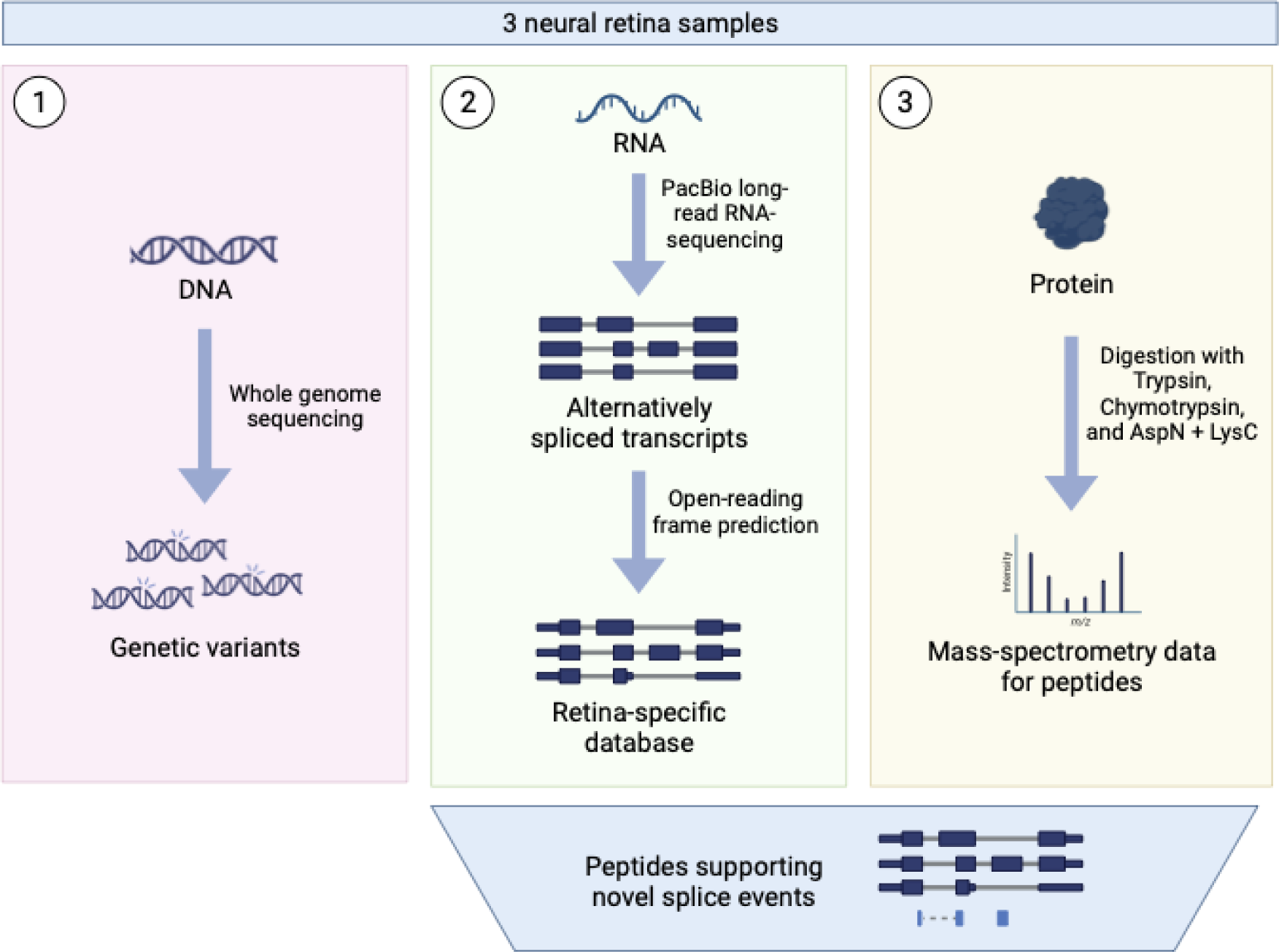

## INTRODUCTION

The neural human retina, located at the back of the eye, is a light sensitive tissue composed of six distinct neuronal cell types (rod and cone photoreceptor cells, bipolar cells, horizontal cells, ganglion cells, and amacrine cells) and Müller glial cells. Its primary function is to convert light into electric stimuli that can be interpreted by the brain. Variants in genes responsible for retina function can disrupt this light conversion process resulting in inherited retinal diseases (IRDs). Approximately 1 in 1,500 individuals worldwide are affected by IRD-associated vision loss (Ben-Yosef, 2022). Similar to other neuronal tissues, the retina is enriched for tissue-specific splicing (Cao et al., 2011; Liu and Zack, 2013). Previous research has revealed retina-specific exons, transcript isoforms, and splicing regulators (Ciampi et al., 2022; Jayasinghe et al., 2018; Murphy et al., 2016). For instance, recent discoveries revealed that the dominant *CRB1* isoform in photoreceptors is not the canonical *CRB1-A* isoform that is expressed in Müller glial cells but the novel *CRB1-B* isoform (Ray et al., 2020). Moreover, it was discovered that splicing factors MSI1 (Murphy et al., 2016) and PCBP2 (Ling et al., 2020) play a role in rod-specific splicing events. It is estimated that at least 11% of causative variants in IRD-associated genes interfere with pre-mRNA splicing (Bacchi et al., 2014; Khan et al., 2020).

Understanding of the expressed transcript and protein isoforms, as well as knowledge of retina-specific splicing events, is required to correctly classify genetic variants identified in IRD patients (Aísa-Marín et al., 2021; Farkas et al., 2013; Swamy et al., 2020). Several mechanisms by which retina-specific splicing influences variant classification have been revealed. Firstly, variants in *ABCA4* and *BBS8* can lead to photoreceptor-specific pseudo-exon inclusions by activating cryptic splice sites or exonic splice enhancers that are only recognized by the photoreceptor splicing machinery (Albert et al., 2018; Murphy et al., 2015; Riazuddin et al., 2010). Other variants, like the frequent deep intronic Leber congenital amaurosis-associated variant c.2991+1655A>G in *CEP290,* demonstrated a higher pseudo-exon inclusion in retinal organoids compared to lymphocytes and fibroblasts (Den Hollander et al., 2006; Parfitt et al., 2016). Another mechanism through which the retina transcriptome affects variant calling involves retina-specific isoforms. Notably, about 80% of variants in *RPGR* are located in a retina-specific exon in a non-canonical isoform called ORF15 (Liu and Zack, 2013). These examples emphasize the importance of studying tissue-specific splicing when interpreting variants in IRD patients.

Several retinal transcriptome studies have been published (Farkas et al., 2013; Pinelli et al., 2016; Ratnapriya et al., 2019; Ruiz-Ceja et al., 2023; Schumacker et al., 2020). However, the existing literature provides limited evidence on the transcript isoforms expressed in the retina for two main reasons. Firstly, the human neural retina can only be accessed post-mortem, and rapid extraction and sample preservation are crucial to maintaining RNA integrity. Therefore, collecting high-quality neural retina samples for transcriptome studies is challenging. Secondly, current datasets are derived from short-read RNA-sequencing, and inference of full-length transcript isoforms comes with uncertainties (Sarantopoulou et al., 2021). Recent advances in long-read RNA-sequencing allow the sequencing of entire transcripts, providing insight into the expressed transcript isoforms and encoded open reading frames (ORFs). Moreover, earlier studies primarily focused on transcript isoforms, but it is also important to investigate the influence of transcriptome diversity on proteome diversity (Smith and Kelleher, 2013).

To address the lack of transcript and protein isoform data of the retina, we generated a comprehensive retina proteogenomic atlas by combining PacBio long-read mRNA sequencing data of three high-quality human post-mortem neural retina samples with mass spectrometry (MS)-based proteomic data and short-read whole-genome sequencing (WGS) data.

## MATERIAL AND METHODS

### Tissue collection for WGS, PacBio mRNA sequencing and mass-spectrometry

Human neural retinal samples (n=3) from non-visually impaired individuals were obtained from Fondazione Banca degli Occhi del Veneto (Venice, Italy), with written consent from the donor’s next of kin to be used for research purposes in accordance with the tenets of the Declaration of Helsinki. Detailed information about the donors is shown in Table S1. The eyes were enucleated within 2-12 hours after donor’s death and the neural retina extraction was performed as described previously (Niyadurupola et al., 2011; Osborne et al., 2016). Briefly, after retrieval of the donor cornea, the eyeball was cut at the ora serrata; iris, lens, and the vitreous body were removed and the retina carefully detached from the sclera and the retinal pigment epithelium, cutting at the optic nerve head. The retinal samples were transferred into cryovials and snap-frozen in liquid nitrogen. Samples were shipped in dry ice.

### DNA isolation and WGS library preparation

DNA was isolated from frozen retinal tissue with the QIAamp DNA mini kit (QIAGEN, Aarhus, Denmark) following the standard protocol. The samples were sequenced using a 2x 150 base pair (bp) paired-end module on a BGISeq500. The minimal median coverage per genome was 30-fold.

### WGS data analysis

Burrows-Wheeler Aligner (v.0.7814) was used to map the reads to the Human Reference Genome build GRCh38/hg38. Single-nucleotide variants (SNVs), structural variants (SVs), and copy number variants (CNVs) were called with Genome Analysis Toolkit HaplotypeCaller (Broad Institute) (Li and Durbin, 2009), Manta structural variant caller (Chen et al., 2016), and Canvas Copy Number Variant Caller (Roller et al., 2016), respectively. After sequencing, the data was processed with an in-house pipeline for pathogenic variant detection using the Human Reference Genome build GRCh38/hg38 according to the procedure of de Bruijn *et al*. (2023). To ensure that no IRD-associated pathogenic variants were observed in our non-IRD samples, we assessed the WGS data for putative pathogenic variants in currently known IRD-associated genes. In short, all SNVs with allele frequencies higher than 0.5% in gnomAD v.3.1.2 were discarded. We prioritized variants with the predicted effects: stop-loss or gain, start-loss or gain, frameshift, in-frame deletion or insertion. Moreover, we included canonical splice variants, missense variants near splice sites, non-canonical splice variants, and deep intronic and exonic splice variants with a SpliceAI score ≥ 0.2. Next, missense variants with a phyloP score ≥ 2.7, CADD_PHRED score ≥ 15, or Grantham score ≥ 80, and silent variants with a phyloP score ≥ 2.7 or a CADD_PHRED score ≥ 15 were prioritized. For remaining variants that met the criteria, we checked the ACMG/AMP classification using Franklin Genoox Platform (https://franklin.genoox.com). Additionally, ClinVar (Landrum et al., 2018) and LOVD (Fokkema et al., 2011) provided information about previous occurrences of the variants in individuals with IRD.

For CNVs and SVs, we prioritized variants with an allele frequency in the inhouse control SV and CNV database smaller or equal to 0.5%. Variants spanning an exon in an IRD-associated gene were evaluated. Inversion events were only considered when at least one of the breakpoints was located within an IRD-associated gene.

### RNA isolation and PacBio library preparation

To obtain the total RNA from human retina samples, 500 µL of Trizol was added and each sample was homogenized in two rounds using a Tissuelyser kit (QIAGEN, Aarhus, Denmark) for 30 seconds at 30 Hz. After a 5-minute incubation at room temperature (RT), 100 µL chloroform was added, samples were mixed, incubated for 3 minutes at RT, and centrifuged at 12,000 g for 15 minutes. Afterwards, the aqueous phase was mixed with glycogen (5 µg/µl) and 1 volume of isopropanol, and the resulting mixture was incubated at 20°C for 75 minutes and subsequently centrifuged at 12,000 g for 30 minutes at 4°C. The supernatant was discarded, and the resulting RNA pellet was further purified and DNAse treated using the Nucleospin RNA Clean-up Kit (Macherey-Nagel, Düren, Germany) according to the manufacturer’s protocol. The total isolated RNA was quantified using a Qubit fluorometer (Thermo Fisher Scientific, Waltham, MA, USA) and RNA integrity number (RIN) values were assessed using a 2100 Bioanalyzer (Agilent Technologies, Santa Clara, CA, USA). The RIN values of the three samples were above 7.0 and 300 ng of RNA input was used to generate the Iso-Seq SMRTbell libraries using the Iso-Seq-Express-Template-Preparation protocol version 2.0 (Pacific Biosciences, California, USA). Libraries were prepared following the standard workflow (for samples composed primarily of transcripts centered ∼2 kb) and using the SMRTbell® library binding kit 2.1 (Pacific Biosciences, California, USA). The on-plate loading concentration of the final Iso-Seq SMRTbell libraries was 80 pM, and a 24-hour movie time was used for sequencing on a Sequel II system (Pacific Biosciences, California, USA).

### Iso-Seq data analysis

The Iso-Seq data was processed as recommended by PacBio. Circular Consensus sequence (CCS) reads with a minimum read accuracy of 0.99, maximum length of 25,000, and minimum number of three passes were generated using CCS (v6.2.0). Lima (v2.4.0) was applied to remove primers and SMRT adapters to generate full-length (FL) reads. Full-length non concatemer (FLNC) reads were generated with Isoseq3 (v3.4.0) refine and converted from BAM to FASTQ. Minimap2 (v2.24) with parameter*s-ax splice-uf--secondary=no -C5 -O6,24 -B4 --MD* was used to align the reads to the Human Reference Genome build GRCh38/hg38. The mapped isoforms were classified using IsoQuant (v.3.1.2) (Prjibelski et al., 2023) with the GENCODE v39 primary assembly annotation (Frankish et al., 2019). As parameters we applied -*-complete_gened, --datatype pacbio_ccs, --fl_data, -- sqanti_output, --count_exons, --model_construction_strategy fl_pacbio, --check_canonical, -- transcript_quantification unique_only, gene_quantification unique_only*, and -- *splice_correction_strategy default_pacbio*.

A list of 294 IRD-associated genes was downloaded from RetNet (Daiger et al., 1998) (19 May 2022) to filter for IRD-associated genes.

### Protein digestions

The neural retinal tissue was homogenated with the PlusOne Sample grinding kit following the standard protocol. Retina protein homogenates were subjected to in-solution digestion using trypsin (Promega), chymotrypsin (Merck), or a combination of LysC (Wako chemicals) and AspN (Merck). For each digest, 10 µg of total protein was reduced by incubation at RT for 20 minutes with 1 µl 10 mM dithiotrytol (Sigma-Aldrich) followed by incubation for 20 minutes at RT in the dark with 1 µl 50 mM chloroacetamide (Sigma-Aldrich). For tryptic digestion, alkylated proteins were pre-digested by addition of 0.2 µg LysC enzyme and incubated for 3 hours at RT. Samples were diluted with 3 volumes of 50 mM ammoniumbicarbonate (Sigma-Aldrich) prior to the addition of 0.2 µg trypsin and subsequent overnight incubation at 37°C. Chymotrypsin digestion was performed by diluting alkylated proteins with 3 volumes of 50 mM ammoniumbicarbonate prior to addition of 0.2 µg chymotrypsin and subsequent overnight incubation at 25°C. For the combined LysC + AspN digestion, alkylated proteins were pre-digested for 3 hours at RT using 0.2 µg LysC after which the sample was diluted by adding 3 volumes of 50 mM ammoniumbicarbonate prior to overnight digestion with AspN at 37°C in 1mM methylamine (Sigma-Aldrich). Digests were diluted 1:1 with 2% trifluoroacetic acid and stored at -20°C prior to shotgun proteomics analysis.

### Shotgun proteomics

All samples were measured using an Evosep One nanoflow liquid chromatography system (Evosep) connected online to a timsTOF Pro2 mass spectrometer (Bruker Daltonics) via a CaptiveSprayer nanoflow electrospray ionization source (Bruker Daltonics). From each digest, 200 ng was loaded onto Evosep tips according to manufacturer’s instructions and separated using the pre-defined 30 samples per day protocol in combination with an Evosep 150 mm x 0.15mm C18 reversed phase column (EV1106 endurance column packed with 1.9 µm C18AQ particles from Dr Maisch) using 0.1% formic acid (Merck) in ultra-pure water (Biosolve) as solvent A and 0.1% formic acid in acetonitrile (Biosolve) as solvent B. The mass spectrometer was operated in positive ionization mode using the instrument default 1.1 second duty cycle time data dependent acquisition parallel accumulation serial fragmentation (dda-PASEF) method. Settings: 100 ms trapped ion mobility spectrometry (TIMS) accumulation and ramp times, 0.6-1.6 1/K0 mobility range, 100 - 1700 m/z range, 10 PASEF frames, 20eV collision energy at 0.6 1/K0 up to 59eV at 1.6 1/K0, dynamic exclusion enabled for 0.4 minutes. The timsTOF Pro2 instrument was calibrated prior to measurements using ESI-Low tune mix (Agilent technologies) infusion.

### Proteomics analysis

To create a retina-specific peptide search database, we modified the long-read proteogenomic pipeline by Miller *et al*. (2022) to the IsoQuant output. We generated Python scripts to convert the IsoQuant output Gene transfer format (GTF) file into a SQANTI-like classification file. The ORFs of novel transcripts were predicted with Coding-Potential Assessment Tool (CPAT) (v3.0.4), and ORFs were called and combined. A GTF file with coding regions was created and the coding sequences were renamed to exons. After using SQANTI3 protein to classify the ORFs, the SQANTI3 classification of the transcripts was replaced by the IsoQuant classification. The untranslated regions (UTRs) were added, proteins were classified, renamed, and filtered for nonsense mediated decay, protein truncation or unlikely protein classification. Philosopher (v4.8.1) was used to add decoys to the PacBio-GENCODE hybrid database. MSFragger (v3.7) was run with open search parameters and digestion enzymes stricttrypsin, chymotrypsin, or aspn and lysc-p with a maximum of two missed cleavages. The fragment mass tolerance was set to 20 ppm. We included carbamidomethyl of cysteine as fixed modification and oxidation of methionine and acetylation of the N-terminus as variable modifications. The false discovery rate (FDR) was 1%, and protein quantification was performed using IonQuant (v1.8.10). We adapted the scripts by Miller *et al*. for the MSFragger output to create files for visualization in the genome browser and to detect novel peptides.

Pyteomics (v4.6) (Goloborodko et al., 2013; Levitsky et al., 2018) was used to perform *in-silico* digestion of the hybrid database into peptides with a minimum length of seven amino acids and a maximum of two missed cleavages. Spectra of novel peptides were visualized with PDV (v1.7.4) for manual validation (Li et al., 2019). The criteria for manual validation included detecting at least eight ions, including both b- and y-ions, a probability higher than 0.95, and the majority of peaks being accounted for by fragment ions.

### Tissue collection for Oxford Nanopore Technology (ONT) sequencing

For the independent ONT validation dataset, rest material from human donors (n=3) without a history or clinical evidence of retinal disease was collected from either Ghent or Antwerp University Hospital tissue banks in accordance with the ethical principles of the Declaration of Helsinki and under ethical approval of the Ethics Committee of the Ghent University Hospital (IRB approval B670201837286). More information about the donors is shown in Table S2. Eyes were transported in CO2 Independent Medium (Gibco) until dissection. Protocols for retina dissection were optimized in-house. To limit the possible effects of autolysis time on RNA integrity, retinas were isolated only from eyes with a total post-mortem interval lower than 20 hours. After visual inspection to exclude any cross-contamination with RPE, neural retinas were immediately processed (total RNA isolation) or snap-frozen and stored at -80°C until used (total RNA isolation).

### RNA isolation and Oxford Nanopore Technology sequencing

Total RNA from post-mortem adult human neural retina was extracted following manufacturer’s guidelines (RNeasy Mini kit®, Qiagen) followed by DNase treatment (ArcticZymes, Tromsø, Norway) and poly-A capture. Poly-A mRNA samples of sufficient quality (RNA Integrity Value, RIN>8.0) were subjected to direct-cDNA (SQK-DCS109, ONT) library preparation following the supplier protocol with minor adaptations. Each library was then loaded (FLO-PRO002, ONT) and sequenced on a PromethION device (ONT) for 72 h. Information about the number of reads can be found in Table S2.

### ONT data analysis (selected genes)

MinKNOW (v5.1.0) was applied to generate fast5 files, which were then base called with Guppy (v6.1.5). Reads were aligned to the Human Reference Genome build GRCh38/hg38 using minimap2 (v2.24) with the -ax splice flags to allow spliced alignments. Alignment files were converted to BAM format, sorted, and indexed using SAMtools (v1.15). StringTie2 (v2.1.1) was run for transcriptome assembly using the -L parameter and the reference transcriptome annotations (Ensembl human release 103) as guide. The three individual GTF files for selected genes were merged with gffcompare (v.0.12.6), and a FASTA file was created using TransDecoder (v.5.5.0). We predicted ORFs using CPAT (v.3.0.4) and added the UTRs to the CPAT output.

### Gene ontology analysis

Gene ontology (GO) enrichment analysis of the 300 most expressed genes was performed using goatools (v1.3.1) and Benjamini-Hochberg false discovery rate correction. We included biological process, molecular function, and cellular component GO terms and used an adjusted p-value cutoff of smaller than 0.05.

### Comparison of PacBio and reference transcripts

To compare the length of PacBio transcripts to reference transcripts, a list of protein coding genes was downloaded from BioMart (14 March 2023) to select protein coding genes from the IsoQuant output. Additionally, a list of protein-coding reference transcripts and their corresponding length was downloaded (12 July 2022).

### Correlation with number of isoforms

The length of the different transcripts for all genes was downloaded from BioMart (16 May 2023) and for each gene, the longest transcript was selected. The FL count for each gene was calculated using the IsoQuant output and the correlation coefficient was calculated using the scipy (v1.10.1) Spearman correlation.

### Enrichment analysis of novel isoforms in RetNet genes

The distribution of known and novel transcripts between all genes and RetNet genes was compared using a chi squared test.

### Length comparison between known and novel transcripts

Comparison of length between novel and known isoforms was tested using a Mann-Whiney-U test.

### CAGE peak analysis

To test if the TSS of the IsoQuant transcripts were supported by CAGE peak data, we made use of data and scripts from SQANTI3 (Tardaguila et al., 2018). We therefore used refTSS CAGE peaks (v3.1) (Abugessaisa et al., 2019) and the search window was set to 50 base pairs upstream of the TSS.

### Comparison to short-read sequencing studies

First, the data of the retina-specific exons from Murphy *et al*. (2016) and Ciampi *et al*. (2022) was converted into a BED file and exons of IsoQuant transcripts were extracted from the IsoQuant GTF file. Bedtools (v2.27.1) intersect with parameters -f, -u and 1 was used to find the previously reported retina-specific exons that were also included in the IsoQuant transcriptome.

### Analysis of novel transcript and protein isoforms

Protein domain predictions were performed with HmmerWeb (v2.41.2) against the Pfam database. AlphaFold (Jumper et al., 2021)predictions were obtained using the Google Colab notebook (v1.5.3) with standard settings. Protein structures were aligned in PyMOL (v2.5.5). The *EPB41L2* brain data and *IMPDH1* exon expression data used for comparison were obtained from the GTEx Portal on 12 June 2023.

## RESULTS

In this study, we created a human neural retina proteogenomic atlas by combining WGS, PacBio long-read RNA-sequencing, and MS-based proteomics data of three healthy human neural retinal samples. DNA-sequencing was used to rule out that the donors were carriers of known pathogenic variants in IRD genes. PacBio long-read RNA-sequencing was applied to discover full length (FL) transcript isoforms in the retina. After ORF prediction on the novel retina transcripts, the MS-based proteomics data was analyzed using a custom peptide search database to identify novel peptides validating novel transcript isoforms. Additionally, selected transcripts were validated using an independent ONT retina dataset.

### Whole-genome sequencing confirms that the donors do not have an IRD-associated genotype

The isolated DNA of the three samples was sequenced with WGS to rule out the presence of known IRD-associated variants among our study participants. We identified a heterozygous frameshift variant c.7027del in *CEP290*, which is classified as ACMG/AMP pathogenic, in sample 2. Since about 20-40% of the population carries a recessive IRD-associated variant, this finding was not unexpected (Hanany et al., 2018). Sample 2 was still included in the dataset because further analysis of *CEP290* did not reveal a second causative allele and we did not observe any *CEP290* transcripts that were only detected in sample2.

### Discovering the transcript landscape in the human neural retina with long-read sequencing

From the three retina samples, we obtained a total of 11.6 M (million) circular consensus (CCS) reads (Table S3) that resulted in 58,541 unique FL transcripts from 15,807 known and 470 novel genes (Table S4). The individual samples had between 3.4 M and 4.2 M CCS reads, which is in line with previous PacBio Iso-Seq studies (Leung et al., 2021; Mehlferber et al., 2022). Sample 1 had the lowest number of reads and in this sample the average CCS read length was 3.3 kb compared to around 2.2 kb for the other two samples (Figure S1A). Detected transcripts had an average transcript length of 3.2 kb, which is longer than the average of 2.4 kb for GENCODE reference transcripts (Figure 1A). For 11,225 genes (68.79%), more than one isoform was identified, and for 4,664 genes (28.64%) five or more isoforms were sequenced (Figure 1B). The number of isoforms was correlated with the longest reference transcript length (Spearman correlation coefficient = 0.19, p-value = 1.92 x 10^-116^, Figure S1B). Furthermore, 213 out of 294 IRD-associated genes extracted from RetNet (72.45%) had more than one isoform, and 99 genes (33.33%) had five or more isoforms (Figure S1C), which is comparable to the results for the whole dataset. The genes with most isoforms were *ATE1* and *TMEM161B-DT*, both with 30 isoforms. Most transcripts (89%) belonged to protein coding genes (Figure 1C) and gene ontology (GO) enrichment analysis of the 300 most expressed genes revealed that these genes are linked to vision (visual perception, phototransduction, visible light) and the nervous system (substantia nigra development, axonogenesis, negative regulation of amyloid-beta formation) (Figure 1D). Moreover, 86% of the identified transcripts were present in at least two samples, indicating that most transcripts are robustly expressed across individuals (Figure 1E). For the transcripts detected only in one sample, 9,313 map to known transcripts and eight to novel transcripts. 141 of these transcripts, all known, belong to IRD genes. Moreover, 73% of single sample transcripts have an FL count of one, and with an average length of 2.0 kb they are shorter than the average transcript.

**Figure 1.**
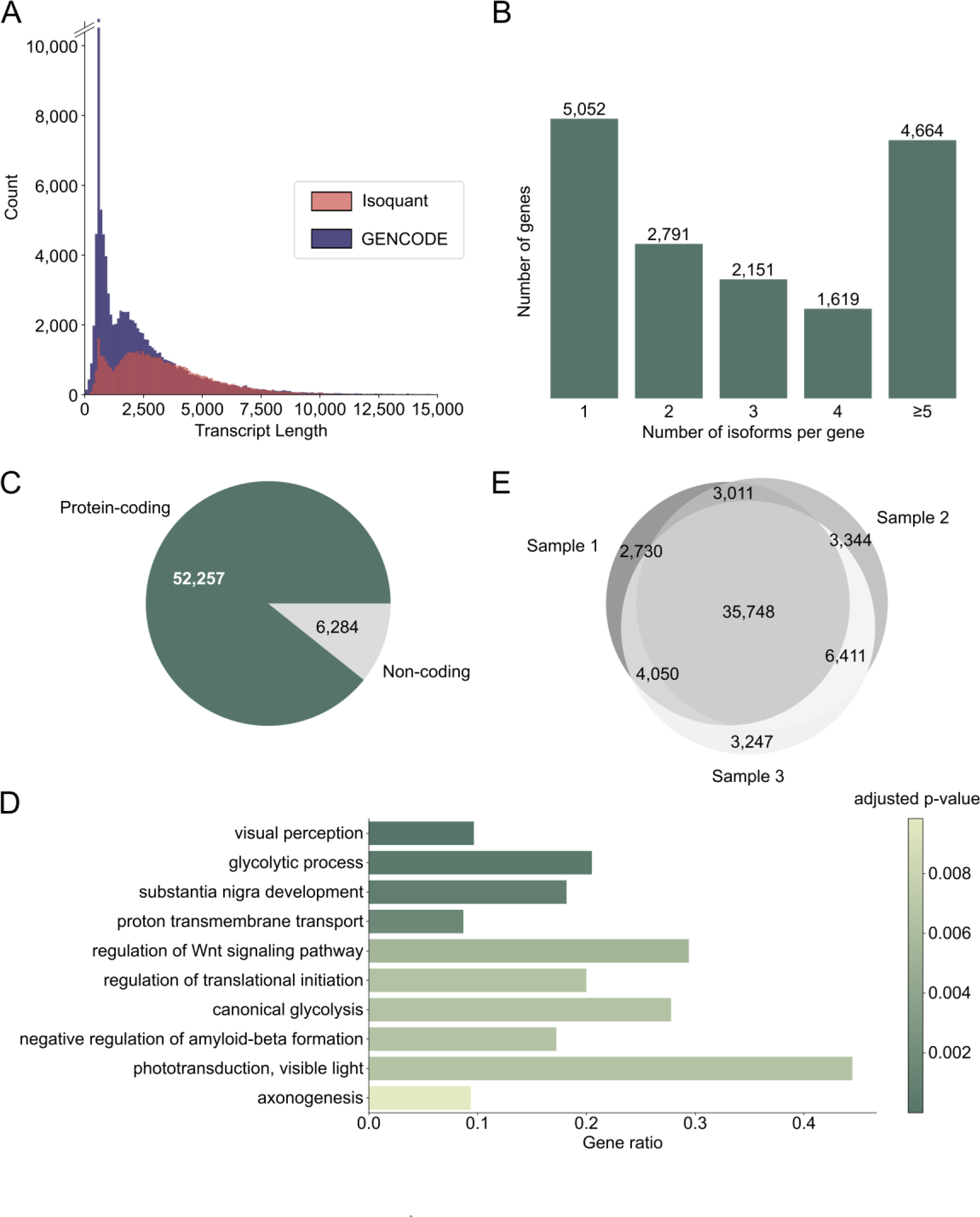
Discovering the transcript landscape in the human neural retina with long-read sequencing. (A) Transcript length distribution of protein coding PacBio IsoQuant (red) and GENCODE reference (blue) isoforms. (B) Number of isoforms detected per gene across the three samples. The number of isoforms for RetNet genes can be found in Figure S1C. (C) Number of isoforms from protein coding genes (green) and non-coding genes (grey). (D) Visualization of the ten most significant hits of the gene ontology (GO) term enrichment analysis of the 300 most expressed genes. The bars are colored by the p-value and the gene ratio shows the percentage of genes associated with the GO-term among the 300 genes. (E) Overlap of transcripts identified in the three individual samples.

### Iso-Seq reveals novel neural retinal transcripts

Following quality control, PacBio transcripts were classified using the IsoQuant classification (Prjibelski et al., 2023). Figure 2A illustrates the different structural categories: Full splice match (FSM), novel in catalog (NIC), and novel not in catalog (NNIC). FSMs match the reference transcript completely, whereas NIC and NNIC are considered novel transcripts. NIC transcripts only contain known splice junctions, however, in new combinations (*e.g.* through the process of exon skipping).

**Figure 2.**
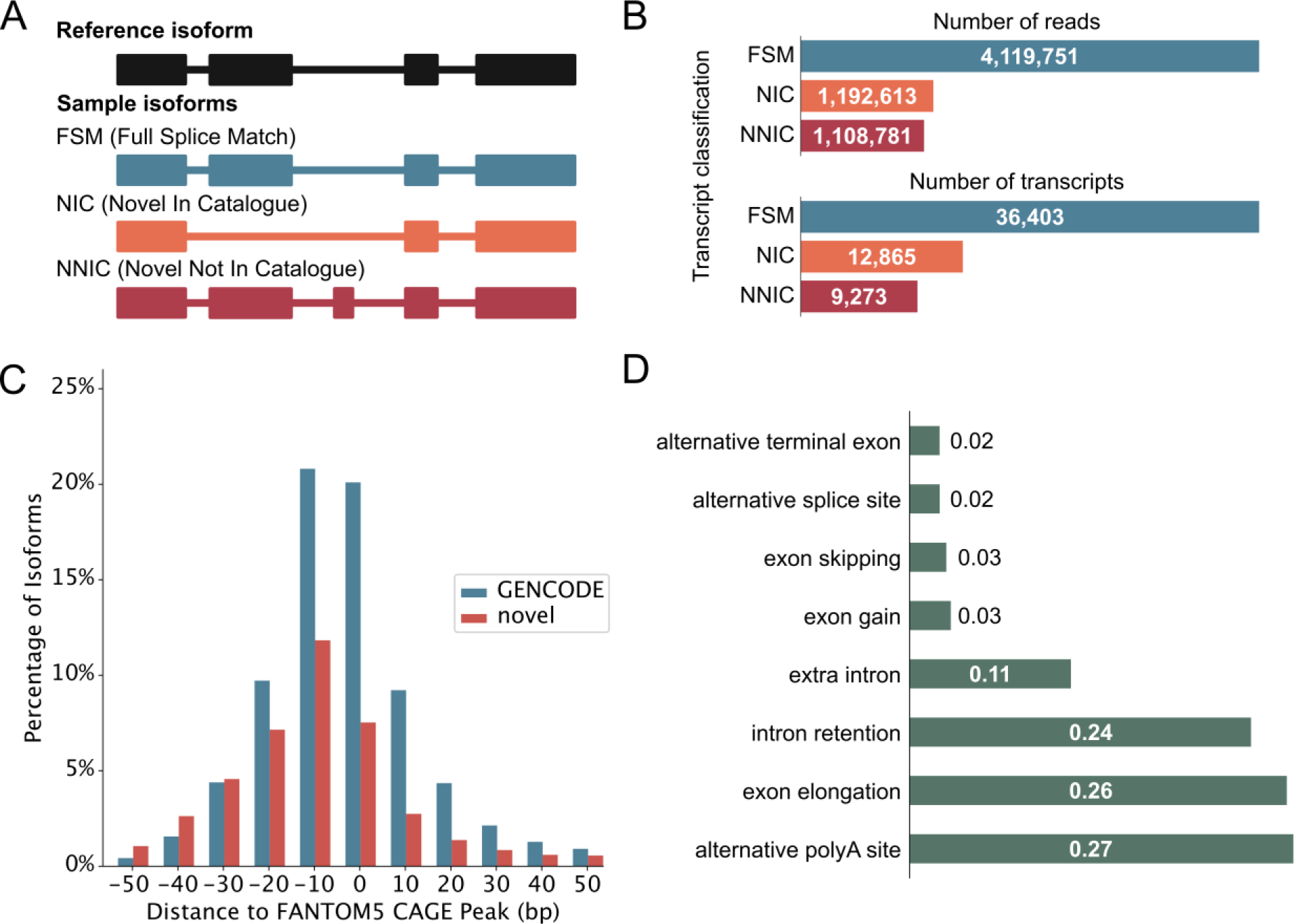
Iso-Seq reveals novel neural retinal transcripts. (A) IsoQuant classification schematic that compares retina transcripts with GENCODE transcripts. Full Splice Matches (FSMs) match the reference completely. Novel transcripts either match the reference splice junctions and are then called Novel In Catalog (NIC), or they contain novel splice junctions and are called Novel Not In Catalog (NNIC). (B) Number of reads and transcripts from the three retina samples associated with each transcript class. The classification of RetNet transcripts is shown in Figure S1D. (C) Distance of transcription start sites (TSS) of known and novel transcripts to annotated refTSS CAGE peaks. (D) Most common novel elements of NIC and NNIC transcripts as classified by IsoQuant. The x-axis represents the fraction of transcripts with that specific event.

NNIC transcripts contain at least one novel splice junction. We identified 36,403 FSMs, 12,865 NICs, and 9,273 NNICs (Figure 2B). The RetNet genes accounted for 522 novel isoforms from 176 genes, and they are enriched in NIC isoforms compared to the complete dataset (X^2^(2, 59741) = 24.99, p =3.74 x 10^-6^) (Figure S1D). Novel transcripts (NIC and NNIC) were longer than known transcripts (Mann-Whitney-U test: W = 5.78 x 10^8^, p-value < -1.80 x 10^-308^) (Figure S1E). To obtain further evidence for the integrity of our novel transcripts, we aligned the 5’-ends of our transcripts with Cap Analysis Gene Expression (CAGE) peak data marking transcription start sites (TSSs). This revealed that 24,382 (66.98%) TSSs of known transcripts and 7,974 (36.02%) TSSs of novel transcripts were overlapping or within 100 base pairs of a CAGE peak (Figure 2C). The most common novel elements in novel transcripts were alternative polyadenylation (poly(A)), followed by exon elongation and intron retention (Figure 2D). Our results are consistent with previous short-read RNA-sequencing studies (Table S5). However, we can call intron retention events with more confidence than in short-read RNA-sequencing studies, because short-read sequencing protocols also capture a fraction of nascent mRNAs that are not yet fully spliced (David et al., 2022). In further support of the quality of our data, we identified eight human homologues out of the ten mice photoreceptor-specific exons identified by Murphy *et al*. (2016) and 45/75 retina-specific short exons and 84/116 retina-specific long exons found by Ciampi *et al*. (2022).

### Most novel ORFs in the IsoQuant-GENCODE hybrid database contain novel protein sequence elements

The long-read proteogenomic pipeline by Miller *et al*. (2022) was used to create a custom peptide sequence database for MS-based proteomic data analysis based on the novel transcripts identified with PacBio sequencing. Our custom database contained 12,499 unique novel ORFs from 22,138 novel transcripts (Table S6). We combined these novel ORFs with 91,124 GENCODE ORFs to form a complete peptide search database (Figure 3A). *In silico* digestion of the ORFs in the hybrid database demonstrated that most theoretical peptides map uniquely to a GENCODE ORF, about a quarter map to both a GENCODE and an IsoQuant ORF, and 1,500 peptides map uniquely to a novel IsoQuant ORF (Figure 3B).

**Figure 3.**
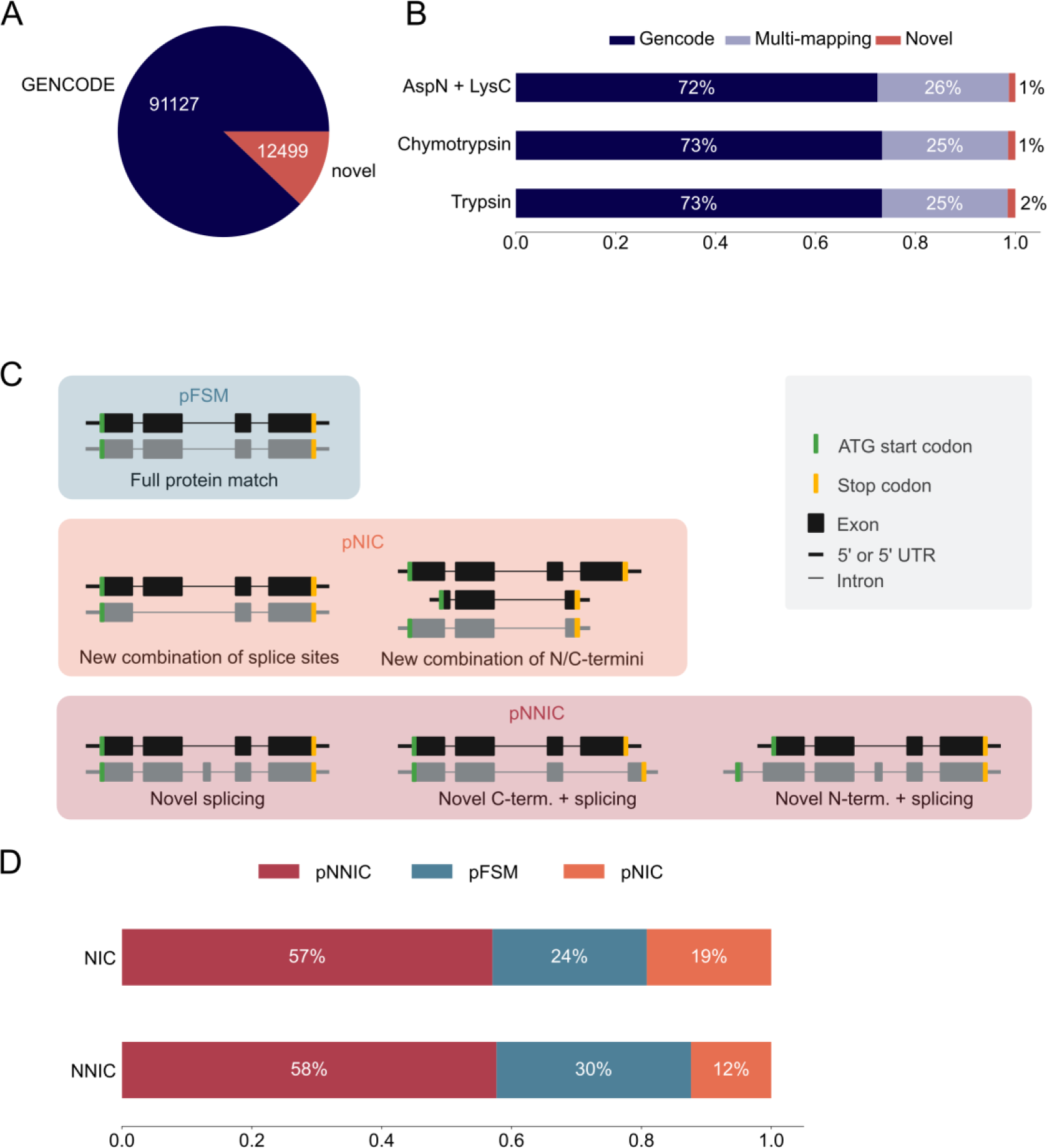
Most novel ORFs in the IsoQuant-GENCODE hybrid database contain novel protein sequence elements. (A) Number of known GENCODE open reading frames (ORFs) (blue) and novel ORFs in the hybrid database (red). (B) *In-silico* digestion of the ORFs in the hybrid database with trypsin, chymotrypsin, or AspN and LysC shows the number of multi-mapping peptides (purple) and uniquely mapping peptides. GENCODE peptides are shown in blue and novel peptides in red. (C) The SQANTI Protein classification scheme compares the long-read ORFs to the reference ORFs. ORFs are classified using the N-terminus, C-terminus, and the splice junctions. A protein full splice match (pFSM) isoform matches a reference isoform. In a protein novel in catalog (pNIC) isoform the N-terminus, the C-terminus, and the splice junctions are known but the combination is new. A protein novel not in catalog (pNNIC) isoform contains at least one novel protein element. (D) Comparison of the transcript (two different bars) and protein (subdivision in the bars) classification of novel ORFs.

Next, we compared the classification of the transcripts with the classification of the ORFs to analyze which transcripts resulted in novel ORFs. Transcripts were classified using IsoQuant as described earlier and ORFs were classified using SQANTI protein (Figure 3C) (Miller et al., 2022). Both classifications are based on the GENCODE reference annotation. A protein full splice match (pFSM) has a known N-terminus, known splice junctions, and a known C-terminus, and the combination of the three protein sequence elements is known. A protein novel in catalog (pNIC) isoform also only contains known protein sequence elements, but the combination of the elements is novel. If at least one of the protein sequence elements, such as the N-or C-terminus or a splice junction, is novel, the protein isoform is classified as protein novel not in catalog (pNNIC). It should be noted that a novel transcript can still encode for a pFSM because the variation can be located in the UTR. We observed that most novel ORFs are classified as pNNIC, followed by pFSM and pNNC (Figure 3D, Table S7). This shows that most novel ORFs are truly novel because of novel protein sequence elements.

### Novel peptides confirm novel IsoQuant isoforms

Digestions with three different enzymes were performed to maximize the sequence coverage of the proteome and the chances of detecting novel peptides. Analysis of the data using the PacBio-GENCODE hybrid database resulted in 33,503 unique peptides from 3,122 unique genes (Table 1). Digestion with trypsin contributed the most peptides (18,671), followed by the combination of AspN and LysC (10,537) and chymotrypsin (8,649). We identified peptides for 15,589 of the GENCODE transcripts and 2,335 of the novel PacBio transcripts. 12,713 peptides were mapped to both a GENCODE and novel transcript while 12 peptides were unique for novel isoforms and 20,723 peptides unique for known isoforms. Ten novel peptides mapping to AMPH, ARHGDIA, EPB41L2, TPM3, and VTI1A passed manual validation (Table 2). The novel peptide in VIT1A initially did not pass manual validation but it was included in the analysis because it was previously detected by Leung *et al* in the cortex (Leung et al., 2021). None of the sequences of the novel peptides were present in UniProt. Alternative splicing events confirmed by novel peptides are exon elongation, novel intron, novel exon, and novel combination of known splice sites. The two peptides that were mapped to intron retention events did not pass manual validation of the spectra. An overview of all novel peptides and their spectra including manual validation is presented in Table S8.

**Table 1:**
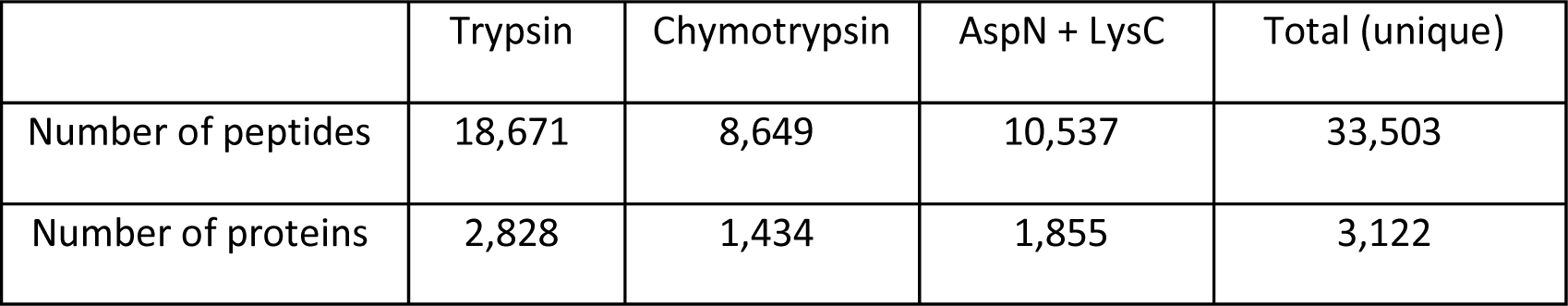
Number of peptides and proteins identified with MSFragger and the custom PacBio-GENCODE hybrid database.

**Table 2:** Overview of the novel peptides detected with the three different digestion enzymes.

| Gene | Digestion enzyme | Transcript | Peptide sequence | Probability<sup>a</sup> | Passes manual validation | Novel event |
| --- | --- | --- | --- | --- | --- | --- |
| *AMPH* | Trypsin | TRANSCRIPT6086.CHR7.NIC | AESSLIEGSER | 0.87 | Yes | Exon elongation |
| *AMPH* | Trypsin | TRANSCRIPT6086.CHR7.NIC | DQDINNSDLSEDEIANQR | 1.00 | Yes | Exon elongation |
| *AMPH* | Trypsin | TRANSCRIPT6086.CHR7.NIC | DTEGLDNSWTHSDVVEHK | 1.00 | Yes | Exon elongation |
| *AMPH* | Trypsin | TRANSCRIPT6086.CHR7.NIC | TLEGTEEFEEK | 1.00 | Yes | Exon elongation |
| *AMPH* | AspN + LysC | TRANSCRIPT6086.CHR7.NIC | DEIANQRYGLLYQEIEA | 0.95 | Yes | Exon elongation |
| *ARHGDIA* | AspN + LysC | TRANSCRIPT33743.CHR17.NNIC | DYMGVGSYSIKSRFT | 0.99 | Yes | Novel intron |
| *ARHGDIA* | AspN + LysC | TRANSCRIPT33743.CHR17.NNIC | TDYMGVGSYSIK | 1.00 | Yes | Novel intron |
| *ATP5PD* | Chymotrypsin | TRANSCRIPT29328.CHR17.NIC | TAQVDAEEKEDVSS | 1.00 | No | Intron retention |
| *CAPG* | Trypsin | TRANSCRIPT16133.CHR2.NIC | MQYAPNTQVRR | 0.81 | No | Intron retention |
| *EPB41L2* | Chymotrypsin | TRANSCRIPT22463.CHR6.NNIC | SHLLESSHETL | 0.99 | Yes | Novel exon |
| *TPM3* | Trypsin | TRANSCRIPT38168.CHR1.NNIC | AISEELDHALNDMTSIASLQPT | 1.0 | Yes | Novel combination of splice sites |
| *VTI1A* | Trypsin | TRANSCRIPT20975.CHR10.NIC | NELLGDDGNSENQLIK | 0.99 | No<sup>b</sup> | Novel exon |

<sup>a</sup>PeptideProphet confidence score
<sup>b</sup>The peptide is still included in the analysis because it was previously identified in an IsoSeq study by Leung *et al.*

Figure 4 shows examples of the novel peptides and their spectra. Five of the novel peptides map to an exon elongation event in *AMPH* (Figure 4A) that is present in three PacBio transcripts (TRANSCRIPT6109.CHR7.NIC, TRANSCRIPT6086.CHR7.NIC, TRANSCRIPT6138.CHR7.NIC), and partly in TRANSCRIPT6110.CHR7.NIC. The transcripts code for proteins with a 102, 406 and 415 amino acid insertion, respectively. The exon elongation could be confirmed by ONT long-read mRNA sequencing on independent retina samples 1-3.

**Figure 4.**
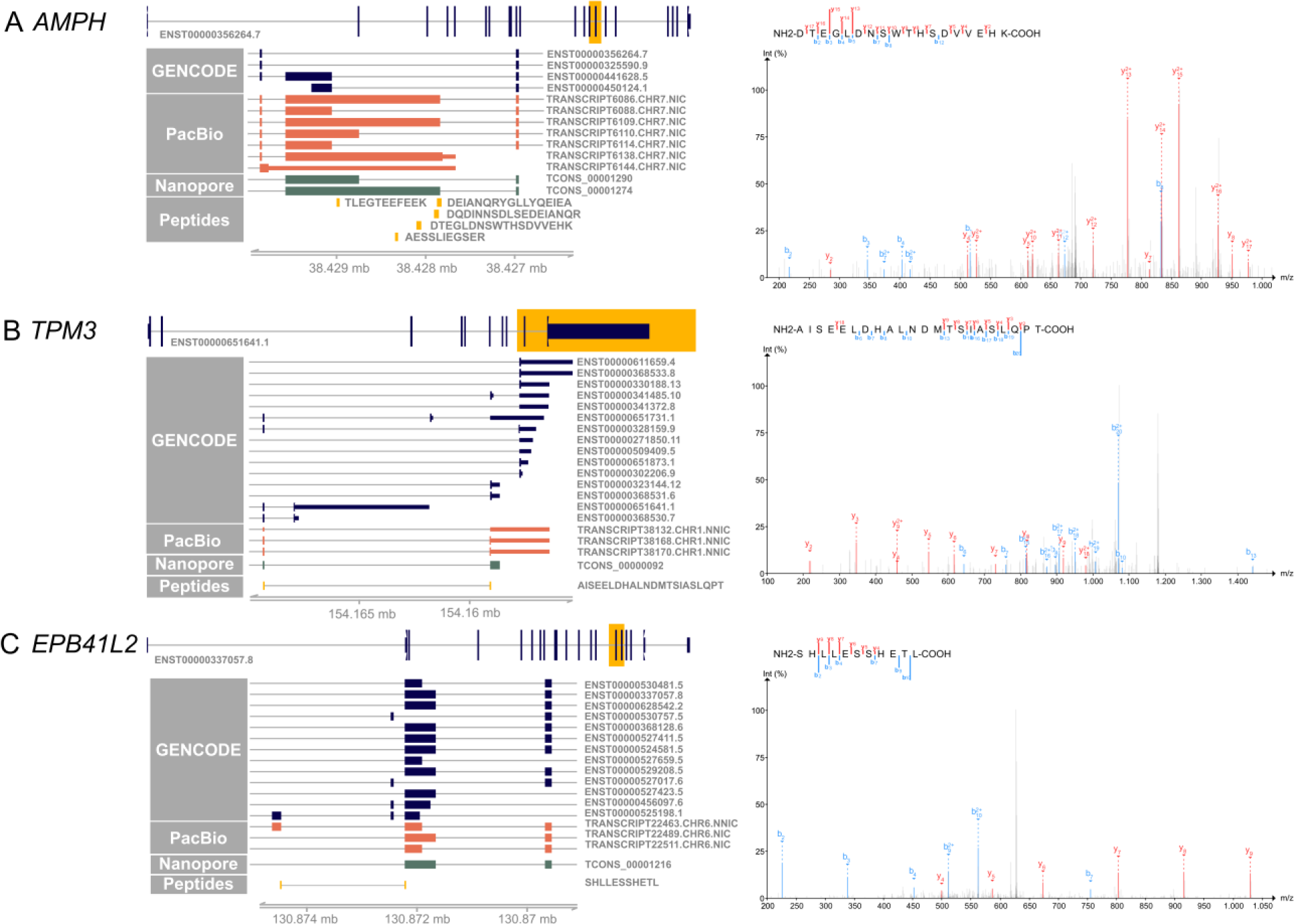
Novel peptides confirm novel IsoQuant isoforms. On the left side, the canonical transcript of each gene is shown with the yellow highlighted region shown in more detail. GENCODE transcripts are shown in blue, novel IsoQuant transcripts in orange, Oxford Nanopore Technology (ONT) sequencing transcripts in green, and novel peptides in yellow. Transcripts were filtered for transcripts with at least one exon in the displayed region, and for the ONT track only one transcript is shown. On the right, the spectrum of the novel peptide is shown with b-ions in blue and y-ions in red. (A) Five novel peptides confirm an intron retention event in *AMPH* that is also supported by the ONT data. The spectrum of one novel peptide is shown, the other four spectra can be found in Table S9. (B) The novel splice peptide in *TPM3* maps to two novel transcripts that contain a novel combination of known splice junctions. (C) A novel splice junction peptide supports TRANSCRIPT22463.CHR6.NNIC derived from the *EPB41L2* gene.

A second example of a novel peptide is AISEELDHALNDMTSIASLQPT in TPM3 that is derived from TRANSCRIPT38170.CHR1.NNIC and/or TRANSCRIPT38168.CHR1.NNIC (Figure 4B). The novel isoforms contain a 79nt penultimate exon that is also observed in reference isoforms ENST00000328159.9 and ENST00000651641.1. Nevertheless, our newly identified isoforms exhibit a distinct final exon compared to the reference isoforms. While they share the last exon with reference isoforms ENST00000368530.7, ENST00000651731.1, ENST00000651641.1, and ENST00000328159.9, the incorporation of the additional exon leads to a reading frame shift and an alternative upstream stop codon. This splice junction is also present in the ONT retina data.

A third example is the novel splice peptide SHLLESSHETL identified in EPB41L2 (Figure 4C). This peptide maps to a novel junction in TRANSCRIPT22463.CHR6.NNIC. The novel *EPB41L2* isoform contains a novel combination of known splice junctions, resulting in an exon coding for 56 amino acids that is also present in two shorter isoforms, ENST00000527017.6 and ENST00000525198.1. In comparison to the two shorter isoforms, the novel isoform does not contain the next small 54 nucleotide exon, like the shorter reference isoforms, but the exon used by most other reference isoforms. The novel junction is not detected in the ONT data but was found with a FL count of 183 in all three PacBio samples.

### RetNet genes demonstrating highest expression of a novel isoform

For 35 out of 294 RetNet genes, the transcript with the novel ORF demonstrated higher expression than all other transcripts from the same gene containing a known ORF. Seven of the 35 genes, *DYNC2H1*, *RPGRIP1*, *KIAA1549*, *LAMA1*, *CEP290*, *DMD*, *USH2A*, and *EYS*, were not completely covered with the PacBio sequencing due to the length of the transcripts (> 8 kb) and were excluded from this analysis. Of the remaining 28 genes, 14 genes had a novel isoform that also contained novel coding sequences (Table 3). Figure 5 shows three examples of these genes.

**Figure 5.**
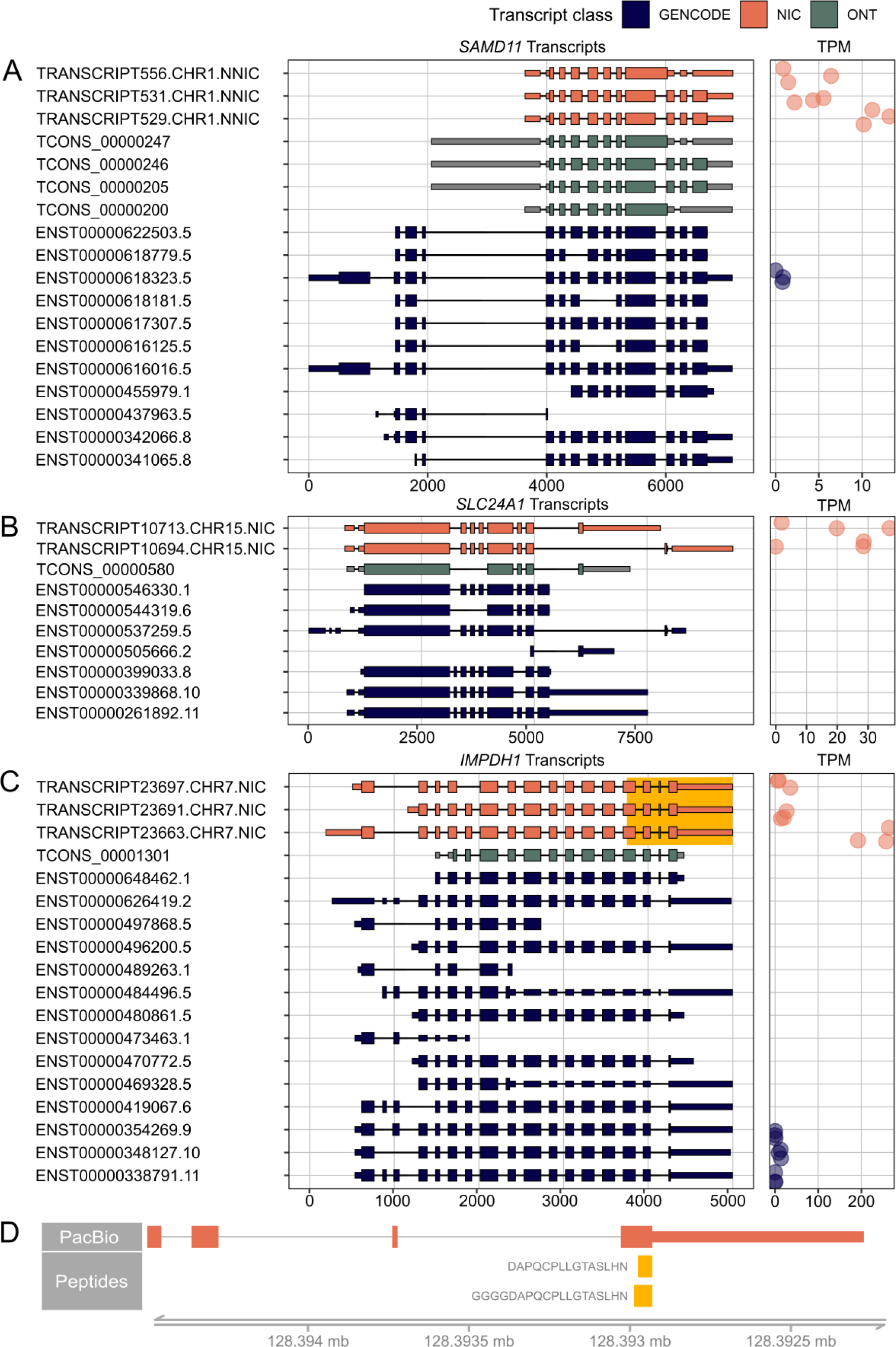
RetNet genes demonstrating the highest expression of a novel isoform. On the left, PacBio transcripts are shown in orange, Oxford Nanopore Technology (ONT) sequencing transcripts in green, and reference GENCODE v39 transcripts in blue. We only show PacBio transcripts that result in a novel open reading frame and ONT transcripts that support novel PacBio transcripts. For all transcripts, the 5’-end is shown on the left and the 3’-end on the right. On the right, the Transcripts Per Million (TPM) count in the three individual samples is shown. (A) *SAMD11* transcripts and their corresponding TPM. (B) *SLC24A1* transcripts and their corresponding TPM. (C) *IMPDH1* transcripts and their corresponding TPM. The highlighted yellow part is shown in (D) with peptides that map to the elongated last exon. The spectra of the peptides are shown in Figure S2.

**Table 3:** RetNet genes with a novel most abundant isoform.

| Transcript | Gene | Length | Structural category | Relative abundance of the isoform | FL count | Subcategory<sup>a</sup> |
| --- | --- | --- | --- | --- | --- | --- |
| TRANSCRIPT4695.CHR10.NIC | *ACBD5* | 5261 | NIC | 0.33 | 158 | Alternative poly(A) site |
| TRANSCRIPT16166.CHR3.NIC | *ATXN7* | 2535 | NIC | 0.31 | 11 | - |
| TRANSCRIPT16650.CHR4.NIC | *BBS7* | 2638 | NIC | 0.54 | 87 | Extra intron, alternative poly(A) site |
| TRANSCRIPT12294.CHR19.NIC | *C3* | 3880 | NIC | 0.88 | 59 | Intron retention, alternative poly(A) site |
| TRANSCRIPT3755.CHR4.NNIC | *CC2D2A* | 5411 | NNIC | 0.17 | 94 | Alternative structure, alternative poly(A) site |
| TRANSCRIPT18031.CHR2.NIC | *CNNM4* | 3830 | NIC | 0.39 | 255 | Alternative poly(A) site |
| TRANSCRIPT47181.CHR1.NIC | *CRB1* | 5341 | NIC | 0.30 | 187 | Intron retention, exon elongation, alternative poly(A) site |
| TRANSCRIPT8849.CHR1.NNIC | *EMC1* | 4913 | NNIC | 1.00 | 267 | Extra intron, intron shift, exon elongation |
| TRANSCRIPT3910.CHR1.NIC | *ESPN* | 6326 | NIC | 1.00 | 565 | Extra intron, exon elongation, alternative poly(A) site |
| TRANSCRIPT9878.CHR13.NNIC | *GRK1* | 7091 | NNIC | 0.65 | 680 | Intron shift, exon elongation, alternative poly(A) site |
| TRANSCRIPT24253.CHR5.NNIC | *GRM6* | 6192 | NNIC | 0.77 | 128 | Extra intron, alternative poly(A) site |
| TRANSCRIPT23663.CHR7.NIC | *IMPDH1* | 2866 | NIC | 0.79 | 867 | Intron shift |
| TRANSCRIPT11816.CHR9.NIC | *INVS* | 7086 | NIC | 0.58 | 90 | Intron retention, exon elongation, alternative poly(A) site |
| TRANSCRIPT20055.CHR11.NIC | *MYO7A* | 10907 | NIC | 0.72 | 76 | Alternative poly(A) site |
| TRANSCRIPT3636.CHR1.NIC | *NPHP4* | 2668 | NIC | 0.39 | 14 | Intron retention |
| TRANSCRIPT1308.CHRX.NIC | *OFD1* | 2539 | NIC | 0.40 | 21 | Intron retention, exon elongation, alternative poly(A) site |
| TRANSCRIPT8654.CHR10.NIC | *PCDH15* | 4743 | NIC | 0.18 | 203 | Intron retention, exon elongation, alternative poly(A) site |
| TRANSCRIPT9279.CHR1.NNIC | *PLA2G5* | 4523 | NNIC | 0.24 | 42 | Alternative structure, intron retention |
| TRANSCRIPT18772.CHR12.NIC | *POC1B* | 3460 | NIC | 0.21 | 39 | Intron retention, exon elongation, alternative poly(A) site |
| TRANSCRIPT18871.CHR6.NNIC | *PRDM13* | 5142 | NNIC | 0.52 | 12 | Alternative structure, alternative poly(A) site |
| TRANSCRIPT26925.CHR17.NIC | *RGS9* | 1667 | NIC | 0.37 | 47 | Extra intron, exon elongation, alternative poly(A) site |
| TRANSCRIPT14425.CHR16.NIC | *RPGRIP1L* | 2528 | NIC | 0.23 | 5 | Intron retention, exon elongation, alternative poly(A) site |
| TRANSCRIPT529.CHR1.NNIC | *SAMD11* | 2382 | NNIC | 0.62 | 42 | Alternative structure |
| TRANSCRIPT10713.CHR15.NIC | *SLC24A1* | 5140 | NIC | 0.50 | 72 | Extra intron, alternative poly(A) site |
| TRANSCRIPT4995.CHR3.NIC | *SLC4A7* | 7577 | NIC | 0.51 | 632 | Exon elongation |
| TRANSCRIPT9410.CHR8.NNIC | *TTPA* | 2466 | NNIC | 0.44 | 11 | Alternative structure, exon elongation, alternative poly(A) site |
| TRANSCRIPT11681.CHR2.NNIC | *WDPCP* | 7044 | NNIC | 0.51 | 54 | Alternative structure, alternative poly(A) site |

NNIC = Novel not in catalog, NIC = Novel in catalog, FL count = Transcript count of the full-length isoform
• Only novel subcategories are shown

We identified a novel major isoform in *SAMD11* (TRANSCRIPT529.CHR1.NNIC) (Figure 5A). This isoform contains a new first exon that is also found in TRANSCRIPT556.CHR1.NNIC and TRANSCRIPT531.CHR1.NNIC. The resulting ORF of TRANSCRIPT529.CHR1.NNIC is missing the N-terminal SAND domain but it still contains the C-terminal SAM domain. The difference between TRANSCRIPT529.CHR1.NNIC and TRANSCRIPT531.CHR1.NNIC is that TRANSCRIPT531.CHR1.NNIC has a longer exon 3. TRANSCRIPT556.CHR1.NNIC contains partial exon skipping of exon 3 and intron retention that results in a frameshift and a preliminary stop codon. All three novel *SAMD11* ORFs were also identified in the ONT data. The difference is that that the ONT transcripts contain an additional non-coding 5’UTR exon, except for TCONS_00000200, which differs in the 3’-UTR from TRANSCRIPT556.CHR1.NNIC.

Figure 5B shows two novel *SLC24A1* transcripts, TRANSCRIPT10713.CHR15.NIC and TRANSCRIPT10694.CHR15.NIC, which both account for 50 percent of the *SLC24A1* transcripts. TRANSCRIPT10713.CHR15.NIC results in a novel ORF while TRANSCRIPT10694.CHR15.NIC corresponds to a known ORF. The novel TRANSCRIPT10694.CHR15.NIC isoform contains an alternative C-terminus with a penultimate coding exon of (33 aa) that is also present in the short reference transcript ENST00000505666.2. The same alternative terminal exon was also identified in the validation ONT dataset as transcript TCONS_00000580.

The third example is a novel *IMPDH1* isoform (TRANSCRIPT23663.CHR7.NIC) that accounts for about 80% of all *IMPDH1* transcripts present in the retina samples (Figure 5C). This isoform is a NIC transcript that combines known splice junctions. The novel part is the inclusion of a small 17-nt exon before the last exon, which results in a reading frameshift and ORF elongation with an alternative stop codon. The same 17-nt exon was also identified in the mouse retina (Wang et al., 2024). This additional exon and alternative stop codon are present in the short reference isoform ENST00000648462.1, but this isoform has an alternative 5’ start, while the novel retinal isoform uses the reference start codon. There are also two peptides, DAPQCPLLGTASLHN and GGGGDAPQCPLLGTASLHN, that map to this alternative last exon (Figure 5D, Figure S2). These overlapping peptides were not called with our novelty analysis because they also map to a reference isoform. However, our transcript analysis shows that the novel isoform is likely to be the most common isoform for *IMPDH1* in the human neural retina, and that this transcript is translated into protein. Therefore, the additional coding sequence should be considered when looking for variants in patients with *IMPDH1*-associated vision loss.

Overall, our study created an important reference dataset for future retina and IRD research. All datasets are available in the corresponding repositories and are also available as genome browser tracks.

## DISCUSSION

The comprehensive proteogenomic atlas generated in our study, combining PacBio long-read RNA-sequencing, MS, and whole-genome sequencing, provides a rich dataset for advancing our understanding of retina-specific transcript-and protein isoforms. We provided several examples of novel protein isoforms supported by MS data, and for many more novel transcripts that are potentially coding for novel protein isoforms. These novel protein isoforms may demonstrate (novel) functionalities different from the reference protein. Moreover, also novel transcripts not validated with mass-spectrometry can be relevant for IRD-research and genetic diagnosis; for example, identified (rare) splice junctions can be predictive of alternative splicing induced by a pathogenic variant. Furthermore, PacBio long-read RNA sequencing adds a unique layer of refinement on retina-specific transcript isoforms, providing information on *in cis* or *trans* occurrence of events. Our dataset does not only identify novel isoforms, but it also confirms known retina isoforms that previously have been assembled from short read RNA-sequencing data. Together, these insights can also provide a valuable resource for therapeutic developments in the context of IRDs.

For *AMPH*, a protein associated with the cytoplasmic surface of synaptic vesicles with enriched expression in the retina and the brain, we observed different amino acid insertions downstream of the BAR domain. We could not identify protein domains in these additional sequences. AlphaFold (Jumper et al., 2021) predicts that the proteins with the insertions have similar 3D structures as the reference protein P49418. The inserted amino acids are part of a loop structure that has a low Local Distance Difference Test score. For the shorter insertion, only loops are predicted, whereas the longer insertions are predicted to fold into three alpha helices which can add a currently unknown functionality to the AMPH protein. It is thus likely that the new isoforms generate stable proteins with slightly different functionalities than the reference protein. Other of novel retina isoforms include *TPM3*, *EPB41L2*, *SAMD11*, *SLC24A1* and *IMPDH1*. The novel *TPM3* isoform contains a 79nt exon inclusion that results in a frameshift and a preliminary stop codon. The alternative exon and corresponding frameshift do not affect any known protein domains.

*EPB41L2* was previously found to interact with cyclic-nucleotide gated channels in the rod outer segments (Cheng and Molday, 2013). The novel isoform accounts for only 4 percent of the *EPB41L2* transcripts, however, it is still detected on protein level. This shows that even low abundance isoforms can be translated into proteins and likely impact the function of the protein. The sequence encoded by the additional exon does not fall into a known protein domain. The canonical transcript ENST00000337057.8 only has a FL count of two in our dataset. The difference between this isoform and all other isoforms detected in the retina samples is that it contains a Spectrin and Actin Binding (SAB) domain. This shows that in the retina mainly EPB41L2 transcripts without SAB domain are expressed, which is also observed in the brain (GTEx).

Mutations in *SAMD11* are associated with adult-onset retinitis pigmentosa (Corton et al., 2016). Within photoreceptor cells, SAMD11 serves as a component of PRC1, required for establishing the proper identity of rod photoreceptors (Kubo et al., 2021). Except for ENST00000618323.5 all *SAMD11* isoforms detected in the retina samples lack the SAND DNA-binding domain. This could mean that in the retina, the majority of SAMD11 proteins do not bind to DNA but instead mainly interacts with other proteins or RNA via its SAM domain.

The *SLC24A1* isoform with the novel ORF contains an alternative terminal exon resulting in a shorter sodium/calcium exchanger protein domain compared to most reference isoforms. However, it is only 10 amino acids shorter than the sodium/calcium exchanger protein domain in transcript ENST00000537259.5.

Variants in *IMPDH1* are associated with retinitis pigmentosa type 10. The most common novel isoform detected in the retina contains a 17nt exon before the last exon resulting in a frameshift with an alternative ORF end of 37 amino acids. This alternative sequence does not affect any known protein domains. According to GTEx, the small exon has no high inclusion in any tissue included in that study, suggesting that this proteoform is retina-specific.

Most novel transcripts do not cause big changes in the encoded protein structure. However, this was also expected because the advantage of long-read sequencing is the sequencing of whole isoforms. Therefore, we expected to mainly find transcripts with novel combinations of known junctions. For *SAMD11*, the novel transcripts miss the part coding for the N-terminus of the protein but contain a novel first non-coding exon resulting in a shortened protein. For *EPB41L2*, *SLC24A1,* and *IMPDH1* we observe a fusion of junctions present in prominent reference isoforms with those identified in less prominent, smaller isoforms. For *IMPDH1*, the new combination of splice junctions results in a frameshift and an alternative last exon or ORF end. There is no frameshift in *SLC24A1,* but the novel transcript also contains an alternative terminal exon.

Moreover, many novel transcripts contained an alternative polyA site, exon elongation, or intron retention. While intron retention and exon elongation might affect the ORF, an alternative polyA site by itself does not. However, it is not uncommon to find different 3’UTR lengths for the same gene as 70% of human genes have more than one polyadenylation site (Navarro et al., 2021). It is known that the 3’ untranslated region (UTR) has regulatory functions as its length can affect translation efficiency, stability of the mRNA, and even tissue-specific expression (Navarro et al., 2021; Tanguay and Gallie, 1996).

Using a conservative approach and manual curation, we detected 12 novel peptides. This number is comparable to the 14 and 30 novel peptides identified by Miller *et al*. (2022) and Mehlferber *et al*. (2022) using a similar approach in cultured T- and endothelial cells, respectively. An important difference between our study and the other two is that we used the conservative IsoQuant tool instead of SQANTI3 (Tardaguila et al., 2018). Our database only contained 12,000 novel ORFs while the other two studies included 35,000 and 27,000 novel ORFs respectively. When we used SQANTI3 or TALON (Wyman et al., 2019) to create the PacBio transcriptome, we also identified more novel transcript isoforms, novel ORFs and novel peptides (Table S9). However, we aimed to create an atlas with high-confidence transcript isoforms, and preferred IsoQuant over SQANTI3 and TALON because of the lower level of false positive isoforms (Pardo-Palacios et al., 2023; Prjibelski et al., 2023). Thus, our study used conservative criteria for novel transcript and ORF detection and finds a similar or larger proportion of novel peptides produced from these ORFs compared to the previously mentioned studies.

To increase proteomic coverage, we performed three different digestions, one with trypsin, one with chymotrypsin, and one with a combination of AspN and LysC. As shown before, using multiple enzymes drastically improves proteome coverage (Sinitcyn et al., 2023). A recent study by Sinitcyn *et al*. (2023) applied deep proteome sequencing to cover a large fraction of the human proteome by MS-based proteomics. Apart from using different digestion enzymes, they applied heavy fractionation and different collision modes, where we used a less accessible biomaterial, only one liquid chromatography fractionation scheme, and one collision mode. Another factor that may have reduced our proteomic coverage in comparison to their study is that many proteins relevant to retina-specific functions and IRDs are membrane proteins that are difficult to detect with MS. Moreover, intron retention was one of the most abundant novel events observed in our study and few intron retention events result in stable proteins due to the presence of premature stop codons. Finally, the number of novel peptides that could be detected was low to start with. A lot of novelty lies in the combination of known splice junctions and in UTR variation.

In conclusion, this study creates a comprehensive overview of the transcript and protein isoforms expressed in healthy human neural retina. Moreover, it highlights the need to study tissue-specific transcriptomes in more detail for better understanding of tissue-specific regulation and for finding disease-causing variants. Choosing a representative transcriptome of the tissue of interest affects variant effect prediction and thereby has an influence on providing a genetic diagnosis to patients with an inherited disease (Salz et al., 2023; Taylor et al., 2015). Therefore, we provide our dataset as reference for future retina and IRD research to contribute to missing heritability in IRD patients.

## DATA AVAILABILITY

The PacBio data underlying this article are available in the European Genome-Phenome Archive (EGA) at www.ega-archive.org and can be accessed with accession number EGAD50000000101. The MS data have been deposited to the ProteomeXchange Consortium at www.proteomexchange.org via the PRIDE (Perez-Riverol et al., 2022) partner repository with the dataset identifier PXD045187. To request access, contact the lead contact. USCS genome browser tracks of the analyzed transcriptomics and proteomics data are available at www.genome-euro.ucsc.edu/s/tabeariepe/retina_atlas. All original code has been deposited at GitHub (www.github.com/cmbi/Neural-Retina-Atlas) and is publicly available as of the date of publication.

## ACKNOWLEDGEMENTS

The graphical abstract, Figure 1A, and Figure 3C were created with BioRender.com. We thank the Radboud Technology Center Genomics for their assistance with the PacBio long-read sequencer, and Rob W.J. Collin, Alex Garanto, Alex Hoischen and Lisenka E.L.M. Vissers for valuable advice on long-read RNA sequencing.

