## Supplementary material for "A proteogenomic atlas of the human neural retina": Figure S1

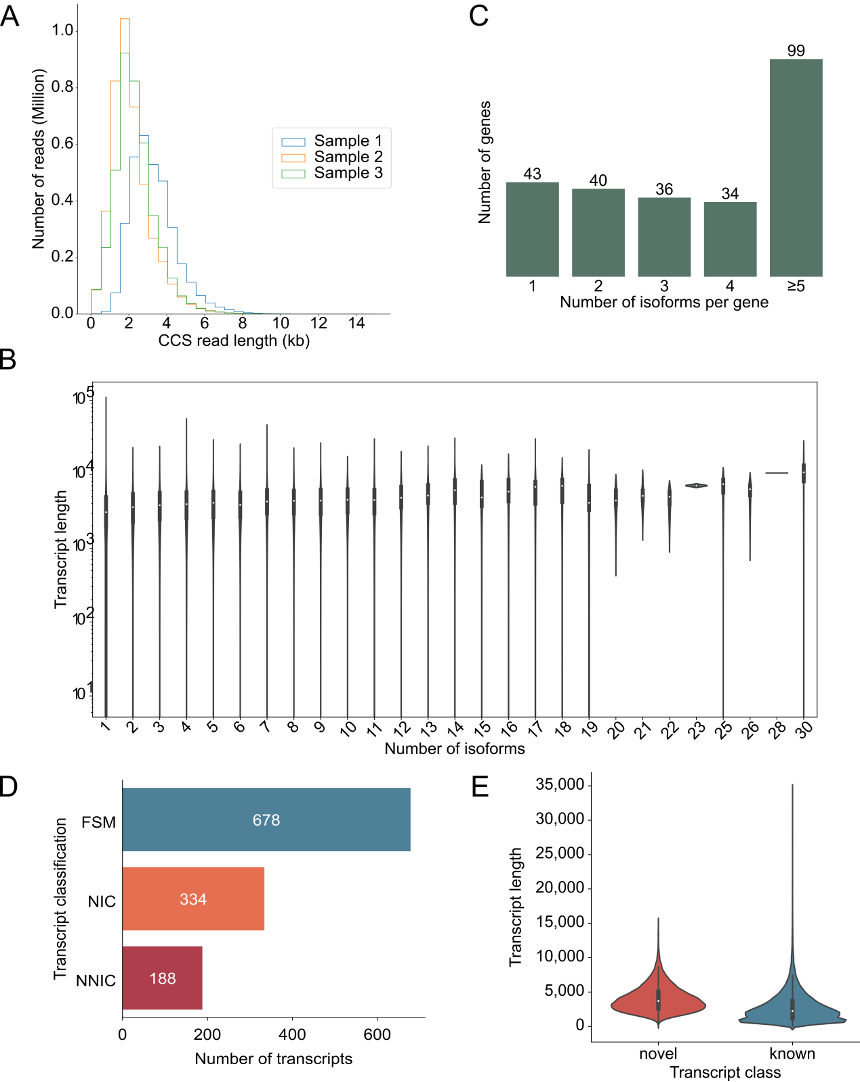

**Figure S1: Additional information about the human neural retina transcriptome generated with PacBio long read sequencing.** (A) Histogram (bin width = 500 bp) of the circular consensus sequence (CCS) read length in retina sample 1 (blue), retina sample 2 (yellow), and retina sample 3 (green). (B) Violin plot showing the gene length depending on the number of isoforms detected for the gene. The Spearman correlation coefficient is 0.19 with a p-value of 1.92 x 10^-116^. (C) Number of isoforms detected per RetNet gene across the three samples. (D) Number of RetNet transcripts from the three retina samples associated with each transcript class. The different classes are Full Splice Matches (FSMs) (blue), Novel In Catalog (NIC) (orange), and Novel Not In Catalog (NNIC) (red). (E) Comparison of the transcripts length between novel transcripts (red) and known transcripts (blue).
