## Supplementary material for "A proteogenomic atlas of the human neural retina": Figure S2

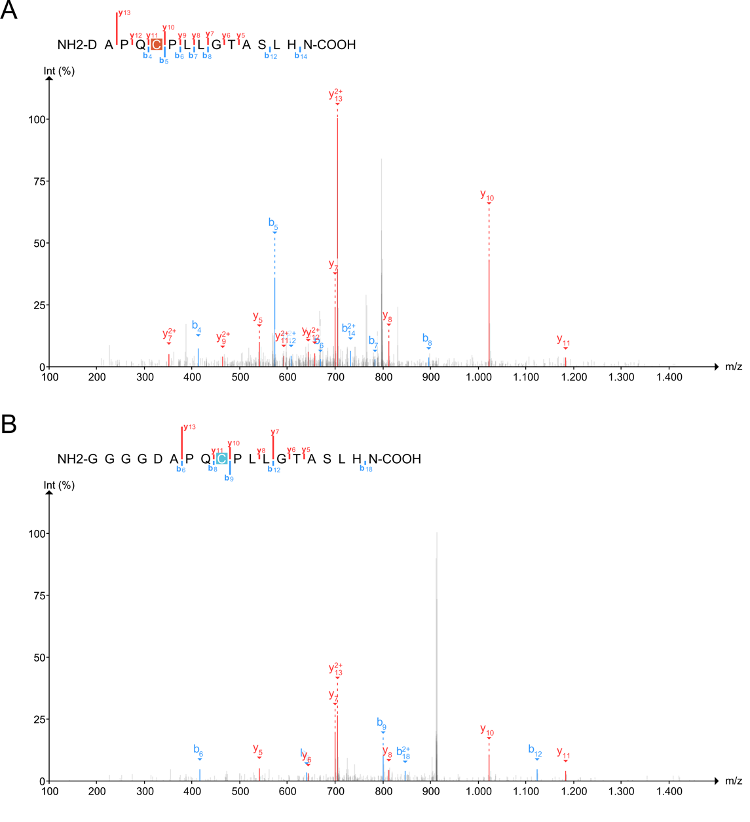

**Figure S2: Spectra of peptides that confirm the alternative terminal exon in *IMPDH1*.** B-ions are shown in blue and y-ions in red.
