## Supplementary material for "A proteogenomic atlas of the human neural retina": Table S1

**Table S1: Description of the samples used for the study.**

| Sample ID | Sex | Age | Time until enucleation | RIN value |
| --- | --- | --- | --- | --- |
| Sample 1 | Male | 59 | 2 hours 35 minutes | 8.2 |
| Sample 2 | Male | 63 | 9 hours 5 minutes | 7.3 |
| Sample 3 | Female | 58 | 11 hours 55 minutes | 7.5 |
