## Supplementary material for "A proteogenomic atlas of the human neural retina": Table S2

**Table S2: Description of the samples used for Oxford Nanopore Technology sequencing.**

|  | Time until enucleation | RIN value | Reads (passed) |
| --- | --- | --- | --- |
| Sample 1 | < 20 hours | > 8 | 10,206,706 |
| Sample 2 | < 20 hours | > 8 | 18,463,865 |
| Sample 3 | < 20 hours | > 8 | 11,388,116 |
