## Supplementary material for "A proteogenomic atlas of the human neural retina": Table S3

**Table S3: Overview of the number of reads for each sample after the different analysis steps.** CCS = circular consensus sequence, FL = full length, FLNC = full length non concatemer

|  | Sample 1 | Sample 2 | Sample 3 |
| --- | --- | --- | --- |
| Reads | 5,063,614 | 7,011,790 | 6,942,704 |
| CSS reads | 3,353,662 | 4,216,550 | 4,044,519 |
| FL reads | 2,956,840 | 3,646,936 | 3,541,082 |
| FLNC reads | 2,949,506 | 3,627,727 | 3,531,995 |
| FLNC with Poly(A) reads | 2,944,441 | 3,616,413 | 3,521,382 |
