## Supplementary material for "A proteogenomic atlas of the human neural retina": Table S7

**Table S7: Comparison of the transcript and protein classification of novel isoforms.** NIC = Novel In Catalog, NNIC = Novel Not In Catalog, pFSM = protein Full Splice Match, pNIC = protein Novel In Catalog, pNNIC = protein Novel Not In Catalog

| Transcript Classification | Protein Classification | Count |
| --- | --- | --- |
| NIC | pFSM | 1883 |
| NIC | pNIC | 1521 |
| NIC | PNNIC | 4526 |
| NNIC | pFSM | 1368 |
| NNIC | pNIC | 565 |
| NNIC | pNNIC | 2636 |
