## Supplementary material for "A proteogenomic atlas of the human neural retina": Table S9

**Table S9: Comparison of the number of transcripts and peptides identified with IsoQuant, SQANTI3, and TALON.**

|  | IsoQuant | SQANTI3 | TALON |
| --- | --- | --- | --- |
| Number of full-length isoforms | 58,541 | 353,381 | 188,346 |
| Number of novel isoforms | 22,138 | 130,796 | 51,477 |
| Number of novel ORFs | 12,499 | 20,330 | 24,892 |
| Number of peptides | 33,503 | 33,539 | 33,450 |
| Number of novel peptides | 12 | 49 | 66 |
